# Resident and elicited macrophages differ in expression of their glycomes and lectins

**DOI:** 10.1101/2020.03.05.975763

**Authors:** Diane D. Park, Jiaxuan Chen, Matthew R. Kudelka, Nan Jia, Carolyn A. Haller, Revanth Kosaraju, Alykhan M. Premji, Melina Galizzi, Alison V. Nairn, Kelley W. Moreman, Richard D. Cummings, Elliot L. Chaikof

**Affiliations:** Department of Surgery, Beth Israel Deaconess Medical Center, Harvard Medical School, Boston, MA 02215; Wyss Institute for Biologically Inspired Engineering, Harvard University, Boston, MA 02115; Department of Biochemistry, Emory University, Atlanta, GA 30322; Complex Carbohydrate Research Center, University of Georgia, Athens, GA 30602

**Keywords:** macrophages, polarization, inflammation, glycosylation, lectins

## Abstract

The pleiotropic functions of macrophages in immune defense, tissue repair, and maintenance of tissue homeostasis are supported by the heterogeneity in macrophage sub-populations that differ both in ontogeny and polarization. Although glycans and lectins are integral to macrophage function, little is known about the factors governing their expression. Here we show that the cellular glycome of murine peritoneal macrophages primarily reflects developmental origin and to a lesser degree, cellular polarization. Resident macrophages were characterized by a simple glycome, predominantly consisting of core 1 O-glycans, while elicited macrophages also expressed core 2 O-glycans, along with highly branched and extended complex-type N-glycans, that exhibited a higher *N*-acetylneuraminic acid:*N*-glycolylneuraminic acid ratio. Strikingly, our analysis revealed that resident and elicited macrophages express 139 lectin genes, with differential expression of 49 lectin genes, including galectins, Siglecs, and C-type lectins. These results suggest that regulation of self-glycan-protein complexes may be central to macrophage residence and recruitment.

## Introduction

Macrophages are specialized leukocytes that help control the recognition and removal of microbes and foreign material, as well as being centrally important in tissue repair and clearance of apoptotic cells. The majority of tissue-resident macrophages originate from mesodermal erythro-myeloid progenitors (EMPs) in the yolk sac and are seeded in the embryo, where they populate tissues and assume distinct phenotypes during organogenesis (*1*). In the peritoneal cavity, two types of macrophages have been described during steady state, large peritoneal macrophages (LPMs), which arise from embryonic precursors, and small peritoneal macrophages (SPMs), which are derived from bone marrow hematopoietic cells (HSCs) (*2*). LPMs represent approximately 90% of resident macrophages and are maintained by self-renewal, independently from terminally differentiated monocytes (*3*). Following induction with inflammatory stimuli, LPMs migrate to the omentum and circulating monocyte precursors infiltrate the peritoneal cavity (*4*). While the mechanisms influencing macrophage population dynamics in various tissues remain unclear, it is generally recognized that tissue-resident and elicited macrophages diverge in development and exhibit functional differences in terms of cytokine secretion, bacterial uptake, and wound healing responses.

In addition to differences in tissue origin and developmental lineage, macrophages also display significant heterogeneity in polarization states. The local milieu, including fragments from apoptotic cells, microbial products and immune signals, can dramatically alter their physiology and functional activity. Macrophages are remarkably versatile and different subsets of macrophages with opposing functions and phenotypes can co-reside in the same tissue. While limitations exist in any classification scheme, a framework has been proposed in which macrophages are categorized into two main subtypes along a continuum: classically activated, pro-inflammatory M1 (LPS and IFN-γ activation) and alternatively activated M2. The M2 subset has been further expanded to include M2a (IL-4 or IL-13 activation), M2b (immune complex and Toll-like receptor activation), and M2c (IL-10 or adenosine activation) populations involved in tissue repair and resolution (*5*). Given such multifaceted roles, macrophages have emerged as therapeutic targets in a number of disease settings, including cancer, autoimmune disease, liver disease, cardiovascular disease, metabolic disease, and inflammatory disease (*6*–*11*). Nonetheless, many unresolved questions remain as to whether switches in macrophage function are associated with discrete alterations in the molecular landscape and whether these alterations can be exploited as disease-related biomarkers or in the assessment of disease resolution.

Traditionally, protein expression has been considered the key phenotype of a cell. Indeed, a number of characteristic proteins have been surveyed and their expression associated with macrophage subsets (*12*–*14*). However, the stability, activity, and function of many proteins are contingent on their post-translational modifications (PTMs). Glycosylation, or the covalent attachment of oligosaccharide chains to amino acids of nascent proteins, is one of the most prevalent types of PTMs. An increasing body of evidence indicates that altered glycosylation is closely associated with disease onset and severity (*15*). Improper glycosylation has been shown to affect processes such as cell differentiation, adhesion, migration, and host protection against bacteria (*16*–*19*). Moreover, glycans that are displayed on the surface in glycoproteins for example are recognized by highly specific receptors, mediating crucial cell-cell interactions. In this regard, the unique repertoire of glycans on macrophage sub-populations may facilitate immune functions and assist in recruiting cells to specific anatomic locations.

Although protein glycosylation plays a critical role in macrophage cell biology, development, and immunology, there is limited information regarding expression of glycans and glycan-binding proteins (lectins) in macrophages according to subtype, thus limiting exploration of their functions. Prior studies surveying the differentiation of the human monocyte THP-1 cell line suggest that glycan changes may occur *in vivo* (*20*, *21*). Here we analyzed the glycans of primary resident and differentially polarized elicited peritoneal macrophages and elucidated key features of glycosylation that are associated with macrophage lineage and polarization. We validated the observed glycan changes by lectin staining and metabolic labeling via cellular O-glycome reporter amplification (CORA) and provide a transcriptomic basis for the altered glycan profiles. In addition, we explored the expression of lectin genes in macrophages, as they are presumed to be key participants in the innate immune system, involved in recognition and removal of pathogens and diseased cells. This study provides new perspective on the interplay between changes in macrophage glycosylation and glycan-binding partners. Subtype-specific glycans may provide targets to modulate macrophage activities, including inflammatory, fibrotic, and reparative responses.

## Results

### Structural elucidation of glycans using mass spectrometry

N- and O-glycans in glycoproteins from distinct macrophage populations, including: resident, elicited (M0), elicited/classically activated (M1), elicited/alternatively activated (M2a, M2c), or elicited/lipid-laden foam cells (**Figure S1**) were released from glycoproteins using PNGase F (N-glycans) and reductive β-elimination (O-glycans), permethylated, and analyzed by high resolution mass spectrometry. Search criteria were established to filter by signal to noise ratio and mass tolerance prior to structural assignment against a reference library (**Figure 1A**; **Table S1**). Matches were validated by tandem mass spectrometry, which provided further evidence of the saccharide compositions and sequence (**Figure S2**). In total, a similar number of compositions were found in all sub-populations (**Figure 1B**). Resident macrophages presented slightly simpler O-glycan compositions than elicited populations. To compare relative abundance levels, quantification was performed by integrating the area per matched glycan, which was then normalized to the sum of all integrated areas in the full mass spectrum.

**Figure 1.**
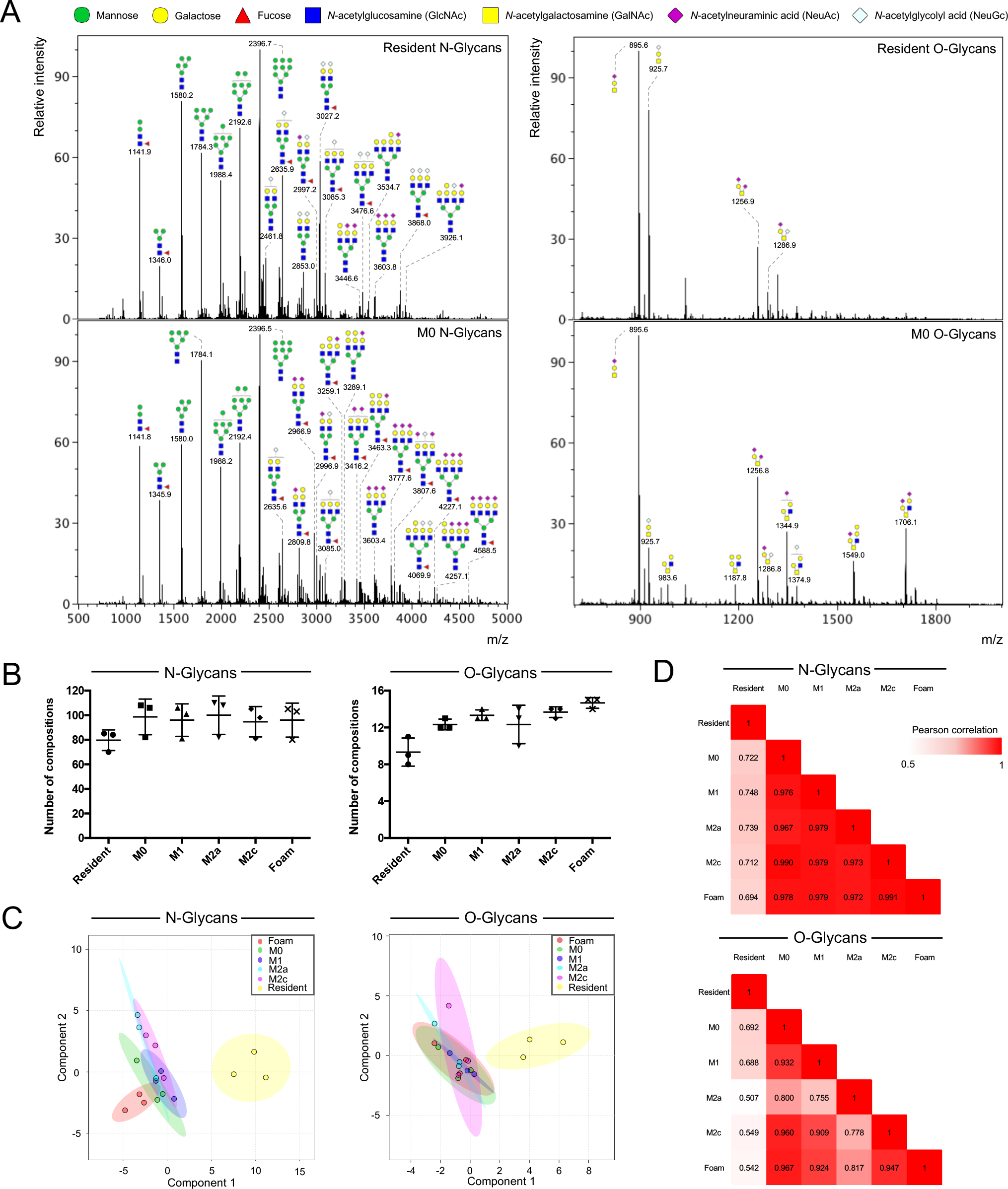
Global N- and O-glycan analysis. (**A**) Representative mass spectra of N- and O-glycans released from resident and elicited/M0 macrophages. Abundant signals are annotated with the corresponding putative glycan structure following symbol nomenclature. (**B**) The total number of glycan compositions identified in each distinct macrophage population. Data are represented as mean ± SD (n=3). (**C**) Partial least squares-discriminant analysis (PLS-DA) score plots for N- and O-glycan profiles of resident, elicited/M0, and elicited/polarized macrophages. The 95% confidence interval is indicated as an ellipse (n=3). (**D**) Correlation matrices of identified N- and O-glycans. Tables are colored according to the numerical values of the Pearson correlation coefficient r between pairs.

Based on the global N and O-glycan profiles, partial least squares-discriminant analysis (PLS-DA) indicated that elicited macrophages are closely related while resident macrophages are distinct (**Figure 1C**). Pairwise Pearson correlation analysis of glycan abundances further supported that resident macrophage glycans deviated the most from those of elicited macrophages (N-glycans, r = 0.722; O-glycans, r = 0.692) (**Figures 1D** and **S3**). Conversely, in elicited populations, M0 glycosylation resembled M1 (N-glycans, r = 0.976; O-glycans, r = 0.932), M2a (N-glycans, r = 0.967; O-glycans, r = 0.800), M2c (N-glycans, r = 0.990; O-glycans, r = 0.960), and foam cell (N-glycans, r = 0.978; O-glycans, r = 0.967) glycosylation. To ensure that similarities in glycan profiles were not a result of inadequate technical processing, we compared the N- and O-glycan profiles of M0 to those of epithelial cells derived from different tissues and found marked differences (HEK293 N-glycans, r = 0.690; O-glycans, r = 0.659; A549 N-glycans, r = 0.772; O-glycans, r = 0.048) (**Figure S3**).

### Comparative N- and O-glycan profiling of resident and elicited macrophage sub-populations

Among all macrophage sub-populations, N-glycan structures were predominantly classified as either high mannose (oligomannose)- or complex-type, with paucimannose- and hybrid-type structures identified as minor components of the total glycome (**Figure 2A**). Complex-type N-glycans exhibited heterogeneity with differentially polarized macrophages expressing a higher proportion of highly branched N-glycans and a lower proportion of biantennary structures than observed in resident macrophages (**Figure 2B**). Specifically, tetraantennary structures were more abundant in M0 (5%), M2a (7%), M2c (6%) macrophages, as well as foam cells (5%) than in resident macrophages (2%). Modification of complex-type N-glycans with fucose and/or sialic acid was common across all macrophage populations, comprising nearly 60% of the total N-glycans observed (**Figure 2C**). Such modified N-glycans bearing both fucose and sialic acid were more abundantly expressed (35-42%) than purely fucosylated (3-5%) or afucosylated structures bearing sialic acid (17-20%) (**Figure 2D**). Among fucosylated N-glycans, mono-fucosylated structures were more common than multi-fucosylated glycans (ratio ~119:1) (**Figure 2E**). In non-humans, sialic acid can occur as either *N*-acetylneuraminic acid (NeuAc) or *N*-glycolylneuraminic acid (NeuGc). N-Glycan structures bearing NeuAc in the absence of NeuGc were among the most abundant species expressed in elicited macrophages, exhibiting a nearly two-fold increase in expression as compared to resident macrophages (**Figure 2F**). Conversely, NeuGc-containing N-glycans, excluding those with both NeuGc and NeuAc, decreased about two-fold.

**Figure 2.**
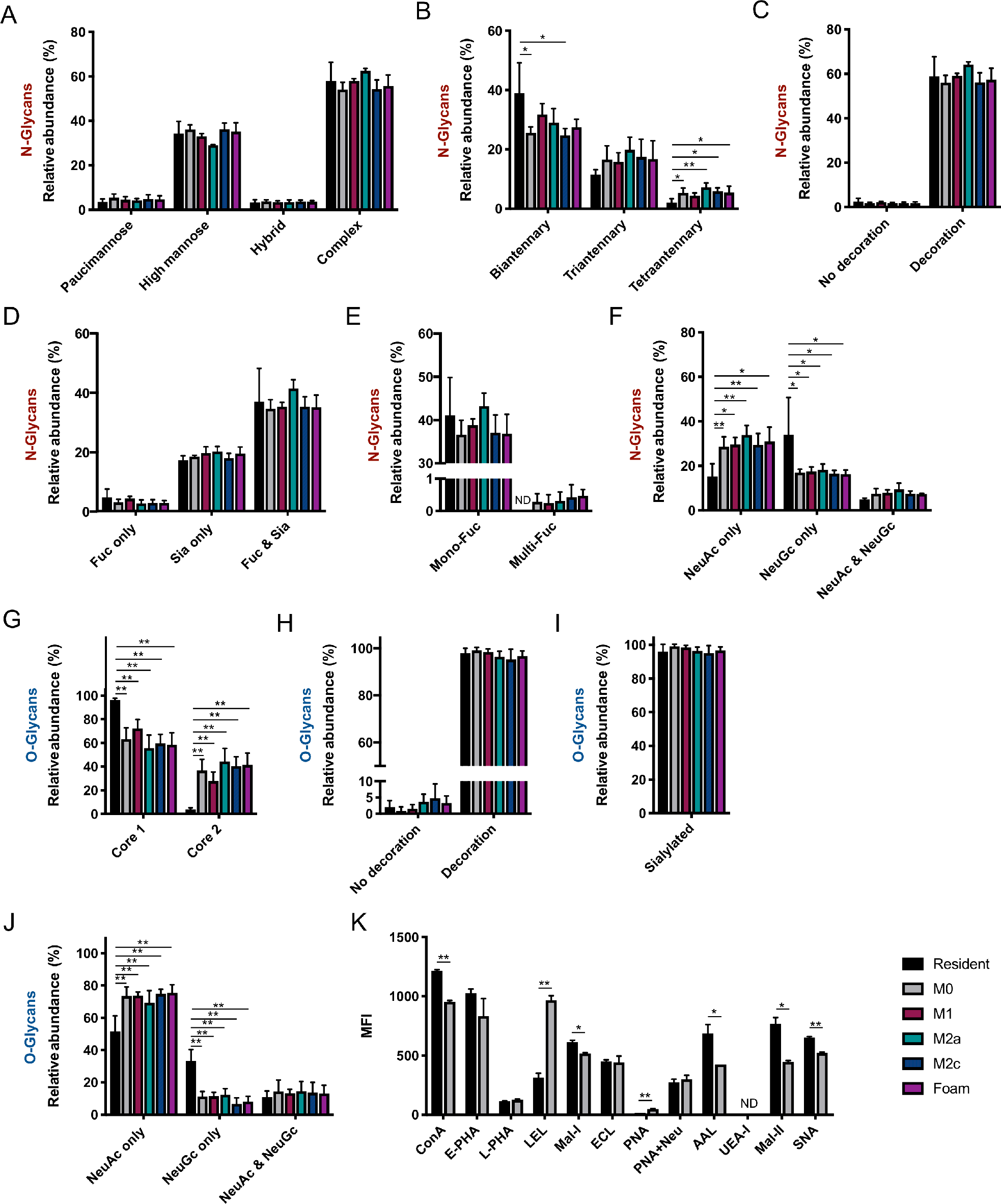
Comparison of N- and O-glycan-specific features among macrophage populations using mass spectrometry and lectin staining. Grouped abundances of N-glycans according to (**A**) class, (**B**) degree of branching, (**C**) presence of decoration, (**D**) type of decoration, (**E**) extent of fucosylation, and (**F**) sialic acid form. Data are represented as mean ± SD (n=3). Grouped abundances of O-glycans according to (**G**) mucin-type core, (**H**) presence of decoration, (**I**) type of decoration, and (**J**) sialic acid form. Data are represented as mean ± SD (n=3). *P*-values were determined using a one-way ANOVA with post-hoc Holm-Sidak comparisons of all pairs relative to resident macrophages. \**P*<0.05; \*\**P*<0.01; Fuc, fucosylated; Sia, sialylated; ND, not detected. (**K**) Lectin staining of resident and elicited/M0 macrophages. Data are represented as mean ± SD (n=2). *P*-values were determined using a Student’s t-test. \**P*<0.05; \*\**P*<0.01; MFI, mean fluorescence intensity; Neu, neuraminidase; ND, not detected.

Mucin-type O-glycans can be classified by their core structures, with resident macrophages predominantly expressing core 1 (96%) along with lower levels of core 2 O-glycans (4%) (**Figure 2G**). Elicited macrophages expressed a higher abundance of core 2-based structures compared to resident macrophages. Among all macrophage sub-populations, the majority (>95%) of O-glycan structures were extended (**Figure 2H**), almost entirely attributed to sialylation (**Figure 2I**). A single mono-fucosylated O-glycan with the composition Hex_1_HexNAc_1_Fuc_1_ (*m/z* 708.34), accounted for the total observed fucosylation and was unique to resident macrophages. NeuAc-bearing structures comprised the majority of sialylated O-glycans, with a modest increase in NeuAc (non-NeuGc) and a pronounced decrease in NeuGc (non-NeuAc) O-glycans observed in elicited macrophages when compared to resident macrophages (**Figure 2J**).

Consistent with these observations, we analyzed surface glycan expression in cells using well-defined lectins known to recognize specific glycan determinants (*22*). Given consistency in all elicited subtypes, lectin binding studies were performed on resident and elicited/M0 macrophages only. Flow cytometry revealed a two-fold decrease in the binding of AAL (binds many fucosylated glycans) and MAL-II lectins (binds sialylated core 1 O-glycans) to elicited macrophages as compared to resident macrophages, due to reduced cell surface expression of α1,6 fucosylation and sialylated core 1, respectively (**Figures 2K** and **S4A-C**). We observed a modest increase in binding of MAL-I (binds α2,3-sialylated type 2 *N*-acetyllactosamine (LacNAc)), SNA (binds α2,6-sialylated type 2 LacNAc), and ConA (binds biantennary N-glycans and terminal mannose on high mannose-type and hybrid-type N-glycans) to elicited macrophages as compared to resident macrophages, suggesting higher levels of galactose (Gal)-β1,4-*N*-acetylglucosamine (GlcNAc), α2,6 sialic acid, and high mannose-type structures, respectively. LEL binding (binds poly-LacNAc (−3Gal-β1,4-GlcNAc-β1-)_n_) was three-fold higher to elicited macrophages than to resident macrophages, indicating greater expression of poly-LacNAc. UEA-1 (binds Fuc-α1,2-Gal-R) did not bind to resident or elicited macrophages, indicating that these cells exhibit little to no expression of α1,2-fucosylation (**Figure S4D**).

In addition to grouped feature analysis, we assessed the significance of glycan compositional changes. Of 118 N-glycan and 15 O-glycan compositions, 11 N-glycans and 6 O-glycans were differentially expressed between resident and elicited macrophages. One of the most abundant O-glycans observed in elicited macrophages, Hex_2_HexNAc_2_NeuAc_2_ (*m/z* 1705.83), was an important distinguishing feature between resident and elicited macrophages (**Figure 3A**). Moreover, this structure was also differentially expressed in M1 macrophages when comparing polarized macrophage sub-populations (**Figure 3B**). Otherwise, consistent with the grouped O-glycan analysis, a core 1-based structure (Hex_1_HexNAc_1_NeuGc_1_; *m/z* 925.43) decreased four-fold in elicited macrophages with a relative shift to sialylated core 2-based structures (*m/z* 1140.55, 1548.75, 1578.76, and 1705.83), which were present in low abundance or not expressed in resident macrophages. Among elicited macrophages, the expression of a single high mannose-type N-glycan, Man_6_GlcNAc_2_ (*m/z* 1783.89), increased nearly two-fold in abundance, while among complex-type N-glycans, a non-decorated structure (*m/z* 1865.94), two mono-NeuGc-containing biantennary structures (*m/z* 2461.22 and 2635.31), and a di-NeuGc-containing triantennary structure (*m/z* 3301.63) decreased or were not expressed, and four NeuAc-containing structures (*m/z* 2156.07, 2966.47, 3776.87, and 3806.88) increased (**Figure 3C**). Two fucosylated and sialylated tetraantennary structures, Hex_8_HexNAc_6_Fuc_1_NeuAc_2_ (*m/z* 4069.02) and Hex_7_HexNAc_6_Fuc_1_NeuAc_3_ (*m/z* 4226.10), were observed in elicited macrophages, but absent in resident macrophages.

**Figure 3.**
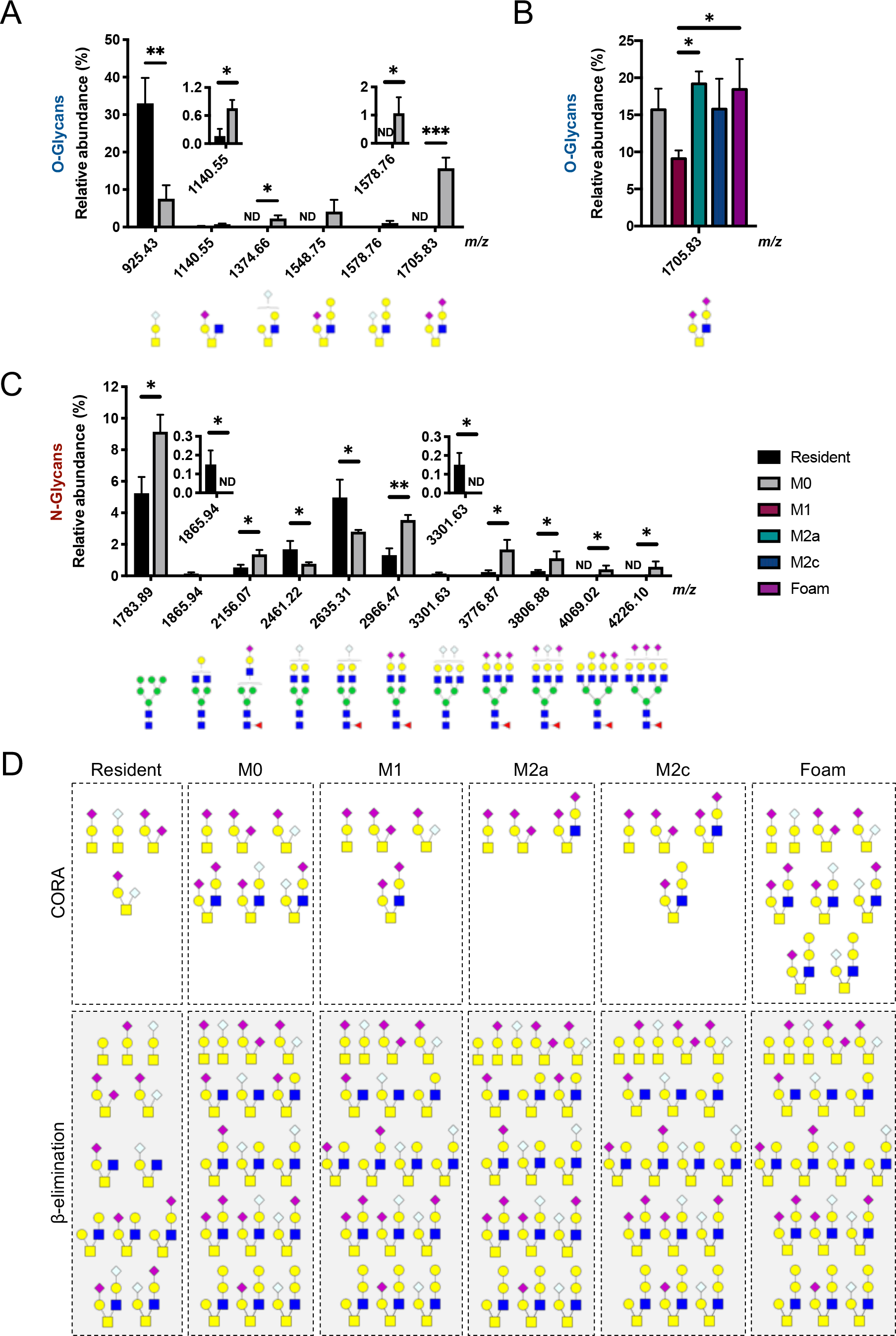
Population-associated alterations of N- and O-glycan compositions. Statistically significant differences between (**A**) resident and elicited/M0 macrophage O-glycans, (**B**) M0 and polarized macrophage O-glycans, and (**C**) resident and M0 macrophage N-glycans. Data are represented as mean ± SD (n=3) and are ordered by increasing mass (left to right). *P*-values were determined using a one-way ANOVA with post-hoc Holm-Sidak comparisons of all pairs. For comparisons where a glycan was detected in only one group, a one sample t-test was used. \**P*<0.05; \*\**P*<0.01; \*\*\**P*<0.001; ND, not detected. (**D**) Putative structures of O-glycans identified in macrophage sub-populations using CORA or β-elimination.

As a complimentary analytical method, and to evaluate the biosynthetic potential of the isolated cells, instead of static glycomic analysis, we performed metabolic labeling of macrophages using cellular O-glycome reporter amplification (CORA). This technique allowed us to examine O-glycans generated by the cells biosynthetically in real time (**Figures 3C** and **S5**; **Table S1**). Core 1-based structures were secreted by all macrophage sub-populations, while core 2-based structures were only produced by elicited macrophages. The CORA results substantially match the results of the O-glycomic analyses, demonstrating that differences in O-glycosylation are derived from biosynthetic pathways operative in the isolated cells.

### Transcriptional basis for altered glycosylation

Glycan assembly requires the coordinated action of glycosyltransferases and glycosidases, which reside within the endoplasmic reticulum (ER) and Golgi apparatus. Of the 148 glycogenes analyzed, 119 were expressed in resident and elicited/M0 macrophages and significant alterations were observed in 45 genes (**Table S2**), of which 40 were associated with glycoprotein and glycosphingolipid pathways and five associated with the synthesis of lipid-linked oligosaccharides (LLO) and glycosylphosphatidylinositol (GPI)-anchored proteins (**Figure 4A**). The differential expression of N- and O-glycan genes between resident and elicited macrophages was further evaluated based on the biosynthetic sequence of respective assembly pathways. Genes participating in the removal of glucose (deglucosylation) of unprocessed high mannose-type glycans (Mogs, Ganab, Prkcsh) were expressed at similar levels in both resident and elicited macrophages, while the expression of Uggt1, responsible for re-glucosylation of Man_9_GlcNAc_2_ in quality control folding of glycoproteins, was elevated six-fold in elicited macrophages (**Figure 4B**). Man1a, which is responsible for the removal of α-linked mannose in high mannose-type glycans that precedes assembly of hybrid- and complex-type structures, was highly expressed in resident macrophages but decreased in elicited macrophages. In addition, multiple genes responsible for the synthesis of complex-type structures and the extension of complex/hybrid-type N-glycans were significantly altered in elicited macrophages. Specifically, Mgat5 and Mgat4a, responsible for β1,6- and β1,4-N-glycan branching, respectively, through the addition of GlcNAc residues, increased and decreased more than three-fold, respectively. Downregulation of Mgat4a in elicited macrophages suggests that the branching GlcNAc residue in triantennary N-glycans is connected by a β1,6- rather than a β1,4-linkage. GlcNAc branches can be further elongated by a galactosyltransferase and expression of the β1,4-galactosyltransferase gene (B4galt3) increased nearly four-fold in elicited macrophages. Expression of Mgat3, which is responsible for the placement of terminal bisecting GlcNAc residues, was four-fold higher in elicited macrophages. Apart from alterations in N-glycan biosynthesis, genes participating in mannose phosphorylation (Gnptab, Gnptg), which target acid hydrolase precursor proteins for transport to the lysosome, were also upregulated in elicited macrophages.

**Figure 4.**
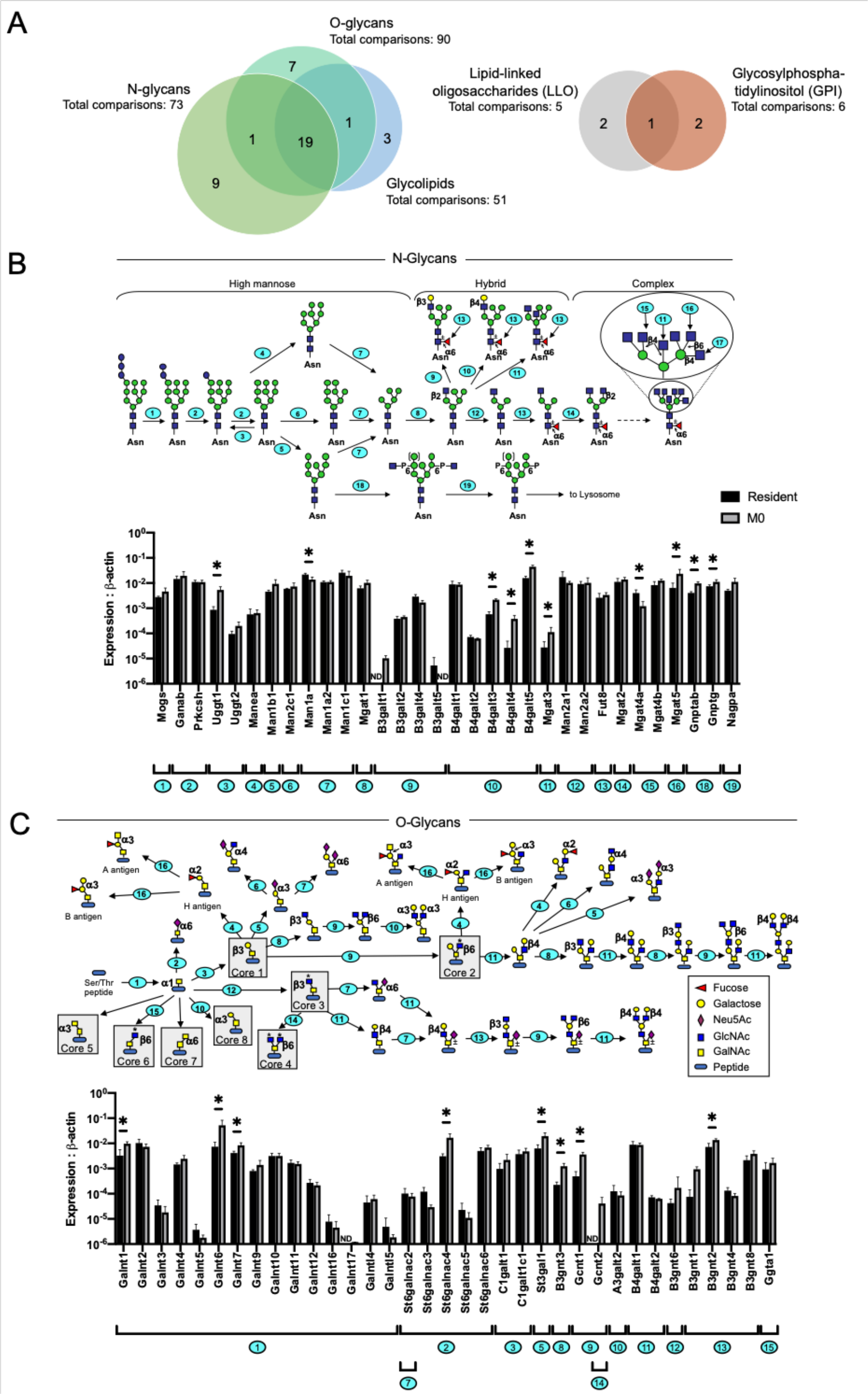
Quantitative RT-PCR analysis of glycosylation-relevant genes. (**A**) Number of genes that showed statistically significant changes between resident and elicited/M0 macrophage populations. Venn diagrams organize genes based on the substrates of the proteins they encode. Biosynthetic pathway of (**B**) N-glycans and O-glycans (top) and the associated changes in the expression of genes involved in each indicated step (bottom), which are numbered in sequence. Data were normalized to the expression of β-actin and are represented as mean ± SD (n=4). \**P*<0.05 (Mann-Whitney U test); -P, phosphorylated; ND, not detected.

While glycan-processing enzymes often recognize both N- and O-glycans, certain glycans are unique acceptors in O-glycan biosynthesis, particularly during initiation. Expression levels of essential genes (Galnt1, Galnt6, Galnt7) involved in the transfer of the initial *N*-acetylgalactosamine (GalNAc) residue to serine or threonine residues of a glycoprotein (GalNAc-α1-Ser/Thr) were significantly higher in elicited than in resident macrophages (**Figure 4C**). Among subsequent biosynthetic steps, the assembly of core 2 O-glycans showed notable differences between resident and elicited macrophages. In particular, Gcnt2 was detected exclusively in elicited, but was absent from resident macrophages, and a closely related gene critical for core 2 formation, Gcnt1, increased more than seven-fold. The GalNAc residue within the core 1 disaccharide Gal-β1,3-GalNAc- α1-Ser/Thr can be modified with sialic acid via an α2,6-linkage and among the *N*-acetylgalactosaminide-α2,6-sialyltransferase genes, enhanced expression of St6galnac4 (five-fold) was observed. Sialic acid capping of terminal galactose residues on N- and O-glycans can also occur via an α2,6- or α2,3-linkage, but among resident and elicited macrophages, α2,6-sialyltransferases were either in low abundance or not detected. In contrast, the α2,3-sialyltransferases St3gal1, which is specific for a Gal-GalNAc moiety, and St3gal6, which is specific for a Gal-GlcNAc moiety, were significantly higher in elicited macrophages (**Figure S6**). Further elongation of glycans by polysialylation was evident based on the expression of St8sia1, St8sia4, and St8sia6 in both resident and elicited macrophages. Of these, St8sia6 decreased 12-fold in elicited macrophages. Collectively, changes among macrophage populations in transcript expression of components of the glycosylation pathways were consistent with glycan structures observed by mass spectrometry and lectin staining (**Table 1**).

**Table 1.**
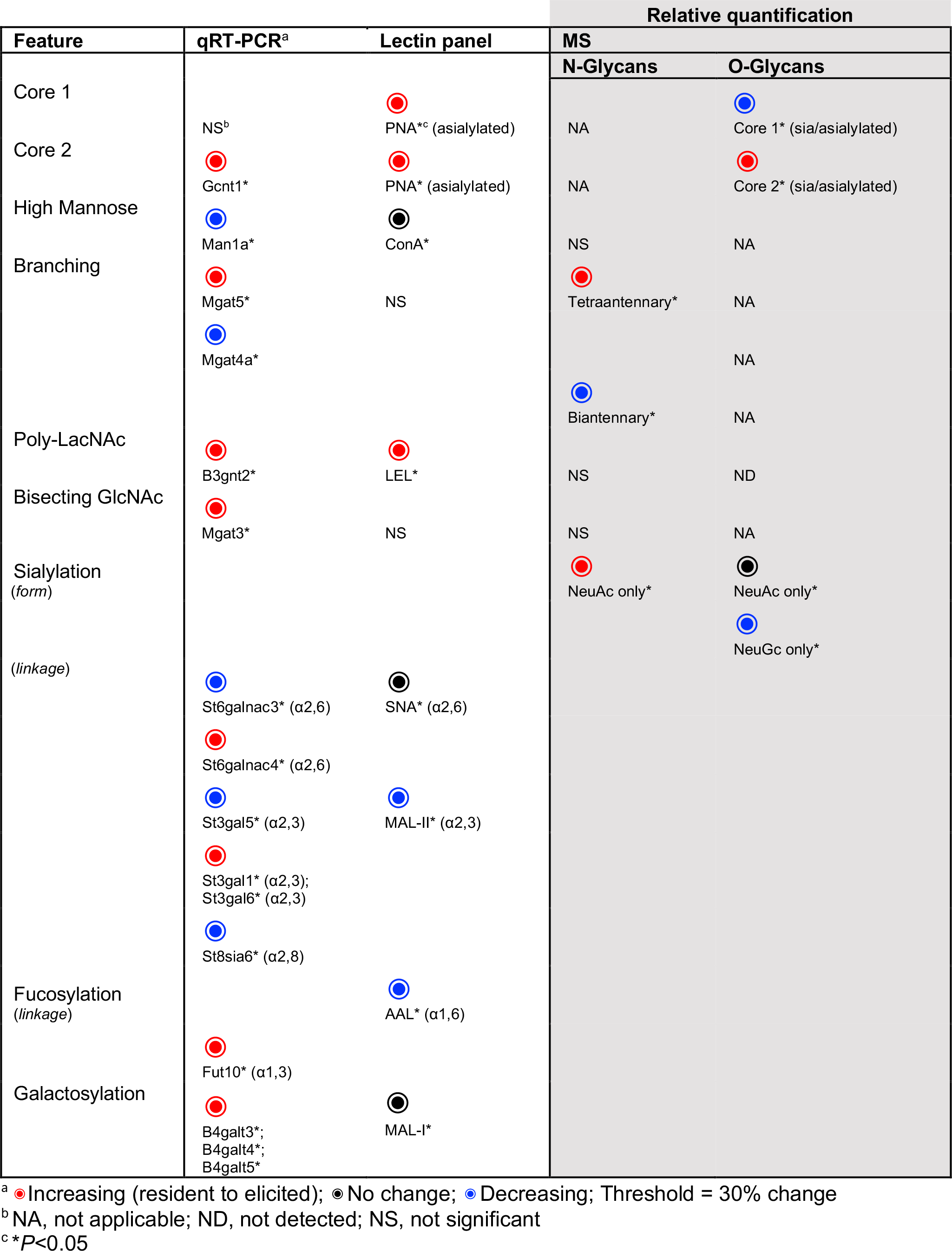
Correlations between glycan feature-related changes as determined by qRT-PCR, lectin staining, and MS analysis.

### Expression of lectins are determinants of specialized macrophage functions

A key enigma in cellular glycomics has been the relationship between the glycans a cell expresses (self-glycans) and the types of lectins it expresses (self-lectins), but this relationship overall has seldom been analyzed. We sought to determine whether the expression of lectins also follows distinct patterns of expression in resident and elicited populations. To this end, we surveyed the expression of 161 known lectin genes with only those genes showing measurable detection levels in at least two of four replicates considered for comparative analysis. In total, we unexpectedly observed that 139 genes were expressed in both or either groups (**Figure S7**; **Table S3**). Enrichment analysis showed that the expressed lectins are predominantly involved in protein processing, phagocytosis, infection, cell adhesion, and cytotoxicity (**Figure 5A**). A total of 49 lectins were differentially expressed, with 38 (78%) relevant to N- and O-glycosylated substrates, five (10%) involved in binding to other glycans, and six (12%) specific to glycolipid recognition (**Figure 5B**).

**Figure 5.**
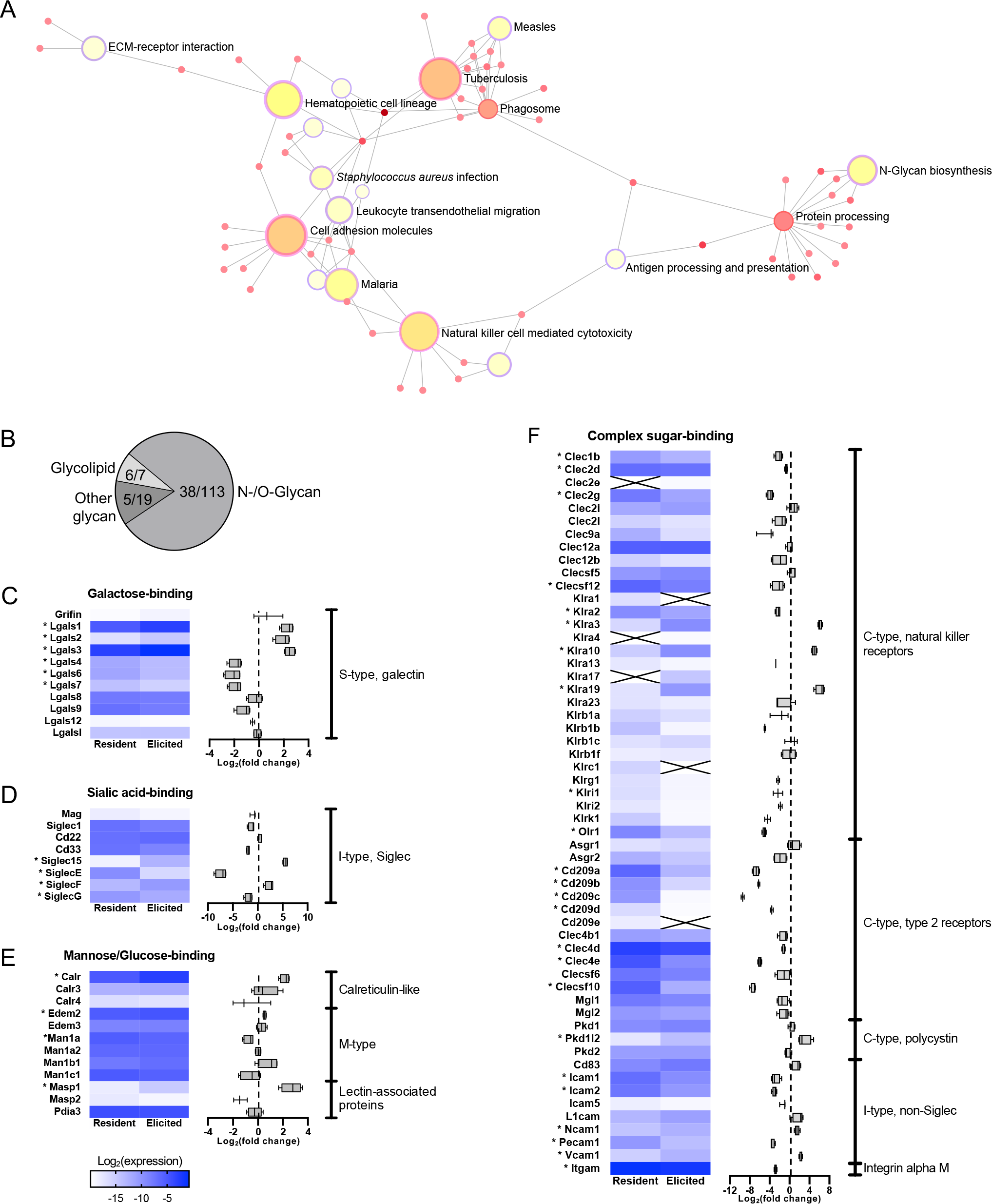
Quantitative RT-PCR analysis of glycan binding protein coding genes. Gene expression in resident and elicited/M0 macrophages were quantified and normalized to the expression of β-actin. (**A**) Bipartite view of enriched gene sets from the KEGG database. Nodes are colored according to *P*-value, sized according to number of genes, and arranged with the Fruchterman-Reingold layout. Nodes with overlapping genes are connected by edges. (**B**) Number of statistically significant genes between resident and elicited macrophages as a fraction of the total number of genes in each group. Genes were grouped by the binding specificity of the protein they encode. (**C-F**) Log_2_ transformed mean relative transcript abundances of N- and O-glycan-specific protein coding genes (n=4). Box and whisker plots represent fold change of gene expression (resident to elicited). Genes within each indicated family are ordered alphabetically. \**P*<0.05 (Mann-Whitney U test); ✕, not detected.

Genes of the galectin (S-type) family that bind to α-galactosides were differentially regulated between resident and elicited macrophages. Notably, Lgals1, Lgals2, and Lgals3 were upregulated, and Lgals4, Lgals6, and Lgals7 downregulated in elicited macrophages (**Figure 5C**). Genes encoding sialic acid-binding proteins termed Siglecs, were also differentially expressed with SiglecG and SiglecE expressed at lower levels, and SiglecF and Siglec15 at higher levels in elicited macrophages, respectively (**Figure 5D**). Mannose- and glucose-specific lectin-coding genes that were differentially expressed were members of the calreticulin-like, M-type, and lectin-associated protein family. The calreticulin-like family of lectins recognize monoglucosylated high mannose-type glycans, which are usually elevated under ER stress or during apoptosis. Calr was abundantly expressed in resident macrophages and increased nearly five-fold in elicited macrophages (**Figure 5E**). M-Type family members, which initiate the degradation of misfolded proteins, showed minimal changes in expression between groups. Unlike other mannose-binding proteins that regulate protein folding, Masp1 and Masp2 encode lectin-associated serine proteases that play a key role in innate and adaptive immune responses. Masp1 increased more than seven-fold in elicited macrophages.

C-Type lectins, which share conserved calcium-dependent carbohydrate binding domains and participate in recognition and initiation of immunity to pathogens, were predominantly downregulated in elicited macrophages. Natural killer cell receptor-coding genes Klrc1 and Klra2 were expressed exclusively in resident macrophages, while expression of Klra17, Klra4, and Clec2e was unique to elicited macrophages. Klra19, Klra10, and Klra3 were expressed in resident macrophages with a significant increase in expression in elicited macrophages. Decreased expression in Olr1, Klrk1, Klrb1b, Klra2, Clecsf12 (Dectin-1), Clec9a, and Clec1b (Clec-2) of more than three-fold was observed in elicited macrophages with a reduction in Olr1 of more than 30-fold (**Figure 5F**). Cd209e (Signr4), which encodes a type 2 receptor, was only detected in resident macrophages. Otherwise, type 2 receptor expression was observed in both resident and elicited macrophages, but typically at reduced levels in elicited macrophages, with particularly pronounced reductions in Cd209b (Signr2) (>650-fold) expression and Clecsf10 (Dectin-2) (>140-fold). Expression of Pkd1l2, a member of the polycystin family, increased 11-fold in elicited macrophages.

Significant differences in expression were also observed for non-Siglec I-type adhesion molecule lectins. For example, expression of Icam1, Icam2, and Pecam1 decreased six-fold, eight-fold, and 11-fold, respectively, in elicited macrophages, while expression of Ncam1 and Vcam1 increased three-fold and five-fold, respectively. The expression of Itgam, which encodes another protein implicated in macrophage adhesion, integrin alpha M, was more than seven-fold lower in elicited macrophages than in resident macrophages. Taken together, these results demonstrate the unique expression patterns for various lectins in resident compared to elicited macrophages, and provide support of an association between self-lectin expression and self-glycans.

## Discussion

Although plasticity is a well-known characteristic of macrophages, the associated molecular changes remain poorly characterized. Based on the hypothesis that glycosylation is cell-type specific and sensitive to environmental changes, we postulated that glycan alterations are an important response that distinguishes macrophage sub-populations from one another. Resident and elicited macrophages were harvested to represent populations of the same anatomical location but with different ontogeny and polarized macrophages were generated from elicited cells to represent sub-populations that have undergone phenotypical changes in response to defined stimuli. By integrating mass spectrometry- and lectin-based approaches to structural glycomics, we generated a map of cell-specific N- and O-glycosylation patterns to identify glycan alterations that may have functional significance. Our analyses revealed that the cellular glycome primarily reflects developmental origin and to a lesser degree the activation state of peritoneal macrophages. Notably, a select set of structures were solely expressed in certain macrophage populations. This analysis demonstrates the value of discerning and monitoring individual glycans, particularly among a pool of structures that share similar features. Conserved glycan signatures across elicited macrophages support the notion that recruited macrophages, as opposed to tissue-resident macrophages, are programmed to respond to acute inflammatory stimuli, which require the participation of glycan-binding proteins. In fact, we found that macrophages express an astonishingly large repertoire of lectins, which is also altered partly upon differentiation. Based on the expression of glycans in macrophage populations, and the specificities of such lectins, we hypothesize that cell surface glycosylation correlates with the specificity and expression of these endogenous lectins. These discoveries highlight the intimate regulation of glycans and their binding proteins in distinct cell populations and provide support that macrophage functions are shaped by cellular ontogeny.

The glycome of elicited macrophages, which are derived from circulating monocytes, differed from that of embryonic progenitor-derived resident macrophages in a number of ways. As compared to resident macrophages, elicited macrophages expressed higher abundances of tetraantennary complex type N-glycans, which was supported by increased expression of Mgat5. The presence of multiple branches provides more sites per glycan for elongation, producing structures that were large in size with diverse compositions. This increase in heterogeneity in elicited macrophage N-glycosylation is likely to affect the cell surface by modulating membrane functions and extracellular interactions. Indeed, N-glycan branching has been reported to enhance the migratory capacity of cells, mediated largely by E-cadherin and integrins (*23*). Furthermore, tetraantennary N-glycans provide increased opportunity for ligand binding and stabilization, which may be required to mount robust responses to growth factors, cytokines, and chemokines during the onset and resolution of inflammation. In fact, Mgat5 deficiency has been shown to diminish surface binding of TGF-β, reduce activation of growth factor pathways, and delay recruitment of leukocytes in murine models of inflammation (*24*). This hypothesis is also consistent with the observed functional differences between resident and elicited macrophages, as elicited macrophages secrete higher levels of pro-inflammatory cytokines and bactericidal reactive nitrogen species in response to LPS or IFN-γ stimuli (*4*). Poly-LacNAc-containing N-glycans were not readily detectable by mass spectrometry, presumably due to their relatively limited expression on subsets of glycoproteins, intrinsic high molecular weight and, as a result, diminished sensitivity in MS. However, lectin staining suggested that a significantly higher number of poly-LacNAc moieties are expressed on elicited macrophages and qPCR confirmed increased expression of B3gnt2 (linear, i-type) and Gcnt2 (branched, I-type), with the latter supporting β1,6-GlcNAc branching as a preferred target for poly-LacNAc extension. Glycan extension by poly-LacNAc may support cellular recognition of selectins and other lectins, as well as other immune regulatory functions (*25*).

Glycan expression patterns that are unique to resident macrophages may be related to their specialized roles within the peritoneal cavity. For example, the high percentage of biantennary N-glycans in resident macrophages may reflect their low proliferative activity, as cells with fewer branched N-glycans exhibit lower surface retention of proteins and display attenuated signaling and growth responses (*19*). Yet resident macrophages express some highly branched structures and are equipped with suitable branching glycosyltransferases. However, unlike elicited macrophages, triantennary N-glycans with β1,4-GlcNAc seem to be dominant in resident macrophages, based on the expression levels of Mgat4a and Mgat5. Interestingly, these structures are associated with functional responses that are more similar to biantennary than triantennary N-glycans with β1,6-GlcNAc linkages (*19*). Peritoneal macrophages are considered to serve as immune sentinels; residing within peritoneal fluid in the immediate vicinity of visceral organs, which they can infiltrate after acute injury (*26*). Although the mechanisms underlying resident macrophage reprograming remain unclear, we speculate that local injury results in the release of metabolic products such as intracellular UDP-GlcNAc and glucose, which increase precursor availability and have been shown to induce N-glycan branching through cellular uptake. Induction of tetraantennary N-glycans on peritoneal resident cells would facilitate proliferation, activation, and tissue repair (*27*).

A prominent difference between resident and elicited macrophages was the presence of significantly greater levels of core 2 O-glycans in elicited macrophages, an observation supported by β-elimination, CORA, lectin staining, and qPCR. Among the diverse functions of O-glycans, core 2 O-glycans have been shown to provide a scaffold for selectin binding ligands (*28*). Loss of core 2 reduces leukocyte rolling activity on immobilized E-, P-, and L-selectin, as well as leukocyte recruitment in thioglycolate-induced peritonitis (*29*). Conversely, the low abundance of core 2 structures in resident macrophages and, specifically, the absence of the disialylated structure, Hex_2_HexNAc_2_NeuAc_2_, may be critical to their ability to maintain tissue homeostasis. Mice engineered with myeloid cells lacking the core 1 synthase specific molecular chaperone (Cosmc) possess the same number of resident peritoneal macrophages as wildtype mice (*30*). However, thioglycolate-elicited macrophages, which are deficient in Cosmc are unable to efficiently clear apoptotic cells.

The display of glycans on cellular surfaces provides specific molecules for recognition by lectins, that can function in both cis- (self) or trans- (non-self) mechanisms. Thus, receptor-glycan interactions initiate a variety of essential intra- and intercellular processes, including cell adhesion, pathogen recognition, cell signaling, and phagocytosis. Interestingly, macrophages express several lectins that can recognize their own glycans in what are referred to as cis interactions (*31*). Galectins are one of the most well studied examples, which can form lattices at the cell surface by binding to galactose residues that extend from cell membrane glycoconjugates. These galectin-glycan clusters are often necessary for regulating cell functions (*32*). Among galectin genes, Lgals3 exhibited the highest fold increase in elicited macrophages when compared to resident macrophages. Galectin-3 (Mac-2 antigen), encoded by Lgals3, shows preference for N-glycans over O-glycans and binds with higher affinity towards tetraantennary chains than less branched structures (*33*, *34*). It has been suggested that the formation of galectin-glycan crosslinks in response to inflammatory stimuli may modulate anti-microbial and anti-apoptotic mechanisms (*35*, *36*), and stabilize cell surface receptors within a molecular lattice to limit glycoprotein endocytosis. Indeed, galectin-3-deficient peritoneal macrophages exhibit reduced resistance to phagocytosis than macrophages from wildtype mice (*37*). Galectin ligation may also regulate T cell activation, as galectin-3 ligation to complex N-glycosylated T cell receptor (TCR) has been shown to reduce TCR sensitivity and suppress T cell activation (*38*). Galectin-1, encoded by Lgals1, displays greater binding to repeating lactosamine units on separate branches, galactose-α2,3-sialic acid, or galactose-α1,2-fucose than to lactosamine units extended in a single chain, galactose-α2,6-sialic acid, or galactose-α1,3-fucose (*33*). Galectin-1 also displays higher avidity toward ligands bound to the cell surface than to those in solution. In this context, higher abundances of tetraantennary N-glycans and increased transcript levels of St3gal1 and St3gal6 in elicited macrophages may account for the concurrent increase in Lgals1 expression, which is known to promote an anti-inflammatory phenotype (*39*).

Sialic acid-binding immunoglobulin-like lectins (Siglecs) are leukocyte lectins that exhibit a wide range of specificities toward sialic acid forms and linkages. While some Siglecs are highly extended (e.g. Siglec-1 and −2) and may interact with trans ligands, many have relatively short lengths, and may typically interact with cis ligands (*40*). We determined that sialylated structures comprise a larger percentage of complex N- and O-glycans on both resident and elicited macrophages than non-sialylated structures. Although the summed abundances of sialylated glycans were comparable in resident and elicited macrophages, differential regulation of Siglecs was clearly observed. Siglec-F and Siglec15, which encode those Siglecs that bind NeuAc-α2,3-Gal and NeuAc-α2,6-GalNAc, respectively, were expressed at higher levels in elicited macrophages than in resident macrophages. This increase may be linked to higher abundances of NeuAc-containing glycans and with the increase in the expression of St3gal1 and St3gal6 in elicited macrophages. Similarly, downregulation of Siglec-G, which binds NeuGc-α2,3-Gal and NeuGc-α2,6-Gal, coincided with a decrease in NeuGc-containing glycans. Siglec-E, which recognizes NeuAc-α2,6-GalNAc and NeuAc-α2,3-Gal, was expressed at significantly higher levels in resident macrophages suggesting that many of the NeuAc-containing O-glycans identified in the resident population are likely linked to GalNAc residues via α2,6- and α2,3-linkages. Interestingly, activated THP-1 monocytes have been shown to express reduced amounts of α2,6-sialic acids (*20*), which contribute to anti-inflammatory responses (*41*). Siglec-E is of particular importance for homeostatic responses as it has been linked to anti-inflammatory signaling via regulation of reactive oxidative species (*42*). Higher expression of Siglec-G in resident macrophages likely also corresponds to their role in regulating B1 cells (*4*).

C-Type lectins serve as non-self (trans) binding receptors and, accordingly, have considerable potential to shape macrophage-mediated responses against microbial and injury related danger signals. Ley et al. have previously shown that C-type lectin expression on macrophages vary depending on the tissue of residence (*43*). We observed that unlike elicited peritoneal macrophages, resident macrophages were characterized by relatively high expression of pattern-recognition receptors (PRRs), including Cd209 (Signr), Clecsf12 (Dectin-1), Clecsf10 (Dectin-2), and Clec4e (Mincle), which recognize cognate microbial carbohydrates. These observations are consistent with the homeostatic and surveillance roles of peritoneal resident macrophages.

Overall we provide a comprehensive analysis of resident and elicited macrophage protein glycosylation, as well as expression of genes regulating glycan biosynthetic pathways and glycan-binding proteins. Our results demonstrate significant differences in macrophage populations, which are consistent with major phenotypic distinctions that contribute to specific macrophage functions. We report this analysis as a resource to examine glycan compositions from primary cells that reflect native glycosylation status, directed toward gaining a full understanding of one of the most structurally complex classes of macromolecules. Capturing molecular details with multiple parallel analytical techniques, as described in this resource, will be useful in deciphering the structural basis of macrophage diversity. The combined broad glycomic profile of peritoneal macrophage populations provides a framework to explore specific glycan and lectin functions and investigate the significance of distinctive glycosylation differences among macrophages that reside in a variety of additional host tissues.

## Materials and Methods

### Macrophage isolation, polarization and characterization

Resident peritoneal macrophages were isolated by lavaging the peritoneal cavity of 8-week-old, male C57BL/6 mice. Elicited peritoneal macrophages were lavaged from the peritoneal cavity 4 days following intraperitoneal injection of thioglycolate. Macrophages were enriched by plate adherence for 1 h and unattached cells were washed off. Macrophages were cultured in complete medium: Dulbecco’s Modified Eagle Media (DMEM) supplemented with 10% fetal bovine serum (Hyclone, GE Healthcare, IL). Where indicated, macrophage activation was induced as follows: 100 U/mL IFN-γ, 100 ng/mL LPS (M1); 20 ng/mL IL-4 (M2a); 100 ng/mL IL-10 (M2c) for 24 h or 100 *μ*g/mL medium oxidized low density lipoprotein (oxLDL) (foam) for 48 h. Because in vitro conditions may affect the cellular glycome, polarization was initiated at different time points so that all cells could be harvested consistently after 72 h of culture. Macrophages were dislocated by non-enzymatic buffer and 5×10^5^ cells were stained at 4°C for 30 min with the following fluorochrome conjugated antibodies: anti-F4/80 (BM8, eBioscience, CA), anti-CD11b (M1/70, eBioscience) anti-I-A/I-E (M5/114.15.2, BD Biosciences, CA), anti-CD86 (GL1, BD Biosciences), and anti-PD-L2 (TY25, BD Biosciences). For staining, cells were cultured in glass slides and polarized to foam cells. Untreated elicited macrophages (M0) were used as a control. Cells were fixed in 10% formalin, washed with water and 60% isopropanol, and let dry. Freshly prepared Oil Red O working solution was added and incubated for 10 min at room temperature. Samples were mounted in anti-fade mounting medium supplemented with DAPI for nuclear staining and imaged by confocal microscopy (Leica SP5 X MP, 63× oil lens, Leica Camera, Wetzlar, Germany).

### N-/O-Glycomic analysis

A total of 1×10^7^ macrophages were collected per cell type to generate both N- and O-linked glycan profiles following previously described protocols (*44*). Briefly, cells were washed with PBS and resuspended in ice-cold lysis buffer (25 mM TRIS, 150 mM NaCl, 5 mM EDTA and 1% CHAPS, pH 7.4). Cell lysis was achieved by sonication on ice. Cell lysates were subsequently dialyzed against ammonium bicarbonate buffer (50 mM) for 48 h at 4°C and lyophilized. The dried glycoproteins were incubated with trypsin for 16 h at 37°C before releasing N-glycans using PNGase F (New England Biolabs, MA). Liberated N-glycans were separated from glycopeptides by C18 Sep Pak SPE cartridges (Waters Corporation, MA). O-Glycans were released from the retaining glycopeptides via reductive β-elimination and purified by co-evaporation with 10% acetic acid in methanol (v/v). Glycans were dried and permethylated prior to mass spectrometric analysis using Bruker Daltonics UltrafleXtreme MALDI TOF-TOF MS (Bruker, MA).

### Cellular O-Glycome Reporter Amplification (CORA)

Resident and M0 macrophages were cultured in complete medium supplemented with 0.1% DMSO or 5 *μ*M Ac_3_GalNAc-α-Bn in 0.1% DMSO, which was generated from GalNAc-α-Bn (Sigma, MO) as previously described (*45*). Media were replaced daily. On day 2, M0 macrophages were polarized to M1, M2a, and M2c for 24 h along with 0.1% DMSO or 5 *μ*M Ac_3_GalNAc-α-Bn in 0.1% DMSO. Conditioned media were collected after 24 h. For foam cell differentiation, M0 macrophages were supplemented with oxLDL on day 1 and 5 *μ*M Ac_3_GalNAc-α-Bn in 0.1% DMSO on day 2. Conditioned media were collected on day 3. Glycans in the media were permethylated and purified prior to MALDI MS analysis as previously described.

### Mass spectrometric analysis

Prepared glycans were solubilized in 10 *μ*L methanol. An aliquot of this suspension (1 *μ*L) was spotted onto a Bruker AnchorChip 384 BC target plate followed by 1 *μ*L of 20 mg/mL 2,5-dihydroxybenzoic acid (Sigma) in 80% methanol, mixed, and dried. Spectra were collected in reflectron positive (RP) mode over a mass range of 700-5000 *m/z* (N-glycans) or 700-2000 *m/z* (O-glycans). Five thousand shots were accumulated per spectrum. Molecular ions were detected as [M+Na]^+^. Mass inaccuracies were corrected using ProteoMass Peptide and Protein MALDI-MS calibration standard (Sigma). Glycan compositions were manually inspected using MALDI LIFT-TOF/TOF MS/MS.

### Data analysis

Glycan compositions were identified according to accurate mass using a library of possible compositions constructed based on the mammalian N- and O-glycan biosynthetic pathways. Signals above a signal-to-noise ratio of 3.5 were filtered and their areas were calculated using flexAnalysis version 3.4 (Bruker). Relative abundances were determined by integrating peak areas for observed glycan masses and normalizing to the summed peak areas of all glycans detected per given sample.

### Lectin staining

Optimal lectin concentrations were determined by titration and specificity was verified by a small molecule inhibitor or pre-treatment with neuraminidase as follows: Peanut agglutinin (PNA) (0.1 *μ*g/mL ± galactose), *Maackia amurensis* lectin I (MAL-I) (5 *μ*g/mL ± lactose), *Erythrina cristagalli* lectin (ECL) (1 *μ*g/mL ± lactose), Concanavalin A (Con A) (5 *μ*g/mL ± mannose), *Aleuria aurantia* lectin (AAL) (1 *μ*g/mL ± fucose), *Ulex europaeus* agglutinin I (UEA-I) (1 *μ*g/mL ± fucose), *Lycopersicon esculentum* lectin (LEL) (5 *μ*g/mL ± chitin hydrolysis), *Phaseolus vulgaris* erythroagglutinin (E-PHA) (1 *μ*g/mL ± thyroglobulin), *Phaseolus vulgaris* leucoagglutinin (L-PHA) (10 *μ*g/mL ± thyroglobulin), *Maackia amurensis* lectin II (MAL-II) (10 *μ*g/mL ± neuraminidase) and *Sambucus nigra* lectin (SNA) (10 *μ*g/mL ± neuraminidase). All lectins were purchased with as biotinylated versions from Vector (Vector, CA). Addition of inhibitors followed manufacturer’s guidelines. Neuraminidase (Roche, IN) pre-treatment of cells proceeded for 1 h at 37°C with gentle shaking every 10 min. A total of 5×10^5^ macrophages were resuspended in 100 *μ*L of staining buffer (1mM CaCl_2_, 1 mM MgCl_2_, 1% bovine serum albumin, 0.1% sodium azide in PBS) per well on a 96-well plate and each well was incubated with one of the 11 biotinylated lectins for 30 min on ice. Cells were subsequently washed twice with staining buffer and incubated with streptavidin-PE (eBioscience) for 30 min at 4°C in the dark. Experimental control conditions included a no staining group and a streptavidin-PE-only group. After further washing, cells were subjected to flow cytometry analysis using BD LSR II (Becton Dickinson, NJ). In addition, cells were stained with only streptavidin-PE as controls to ensure minimal secondary staining. The study was repeated once and an average of replicates is presented.

### RNA isolation and qRT-PCR analysis

Total RNA isolation and cDNA synthesis was carried out with four biological replicates of each cell type as described previously (*46*). The qRT-PCR reactions were performed in technical triplicates for each gene analyzed, which were then averaged. Optimized amplification conditions were applied and data analysis was performed as described previously (*47*). Briefly, Ct values for each gene were normalized to the control gene, β-actin, prior to calculation of relative transcript abundance. In total, 148 glycosylation genes and 161 lectin genes along with 7 reference genes (Cd14, Cd68, Epha2, Adgre4, Mogat1, Mogat2, Sppl3) were included in the analysis.

### Statistical analysis

Partial least squares discriminant analysis (PLS-DA) was performed after autoscaling using MetaboAnalyst version 4.0 (*48*). Statistical evaluation of significant glycan abundance changes was performed using a Student’s t-test. For multiple comparisons, a one-way analysis of variance (ANOVA) was performed followed by a Holm-Sidak test. For comparisons where a glycan was detected in only one group, a one sample t-test was used. Statistical evaluation of significant transcript abundance changes was performed using a Mann-Whitney U test. *P* values of less than 0.05 denote statistical significance. Gene expression was analyzed using NetworkAnalyst 3.0 (*49*).

## Acknowledgments

This work was supported by grants from the NIH, including GM103490 (K.W.M.), HL110843 (A.M.P.), K12 HL141953 (J.C.), P41GM103694 (R.D.C, E.L.C.), and R01DK107405 (E.L.C.). We thank members of the Chaikof and Cummings laboratories for helpful discussions.

## Author Contributions

D.D.P., J.C., M.R.K., C.A.H., R.D.C., and E.L.C. designed research; D.D.P., J.C., M.R.K., N.Y., A.V.N., R.K., and A.M.P. performed research; D.D.P. and M.R.K. analyzed data; A.V.N. and K.W.M. contributed new reagents/analytic tools; D.D.P., J.C., C.A.H., R.D.C., and E.L.C. wrote the paper.

