## Supplementary Material for "Resident and elicited macrophages differ in expression of their glycomes and lectins"

**Supplementary Figures:**

Figure S1. Characterization of murine peritoneal macrophage subsets.

Figure S2. MALDI-TOF/TOF-MS/MS analysis of permethylated N- and O-glycans obtained from peritoneal macrophage subsets.

Figure S3. Correlation scatter plots of N- and O-glycans from M0 vs. resident macrophages, polarized macrophages, HEK293, and A549.

Figure S4. Optimization of lectin staining against elicited macrophages.

Figure S5. MS profiles of permethylated O-glycans in peritoneal macrophage subsets obtained from CORA analysis.

Figure S6. Quantitative RT-PCR analysis of common genes in the N-glycan, O-glycan, and glycolipid biosynthetic pathways.

Figure S7. Quantitative RT-PCR analysis of glycan binding protein genes.

**Supplementary Tables:**

Table S1. N- and O-glycans observed in murine peritoneal macrophage subsets.

Table S2. Relative transcript expression of glycosylation related genes in murine resident and elicited macrophages.

Table S3. Relative transcript expression of lectin genes in murine resident and elicited macrophages.

### Supplementary Figures

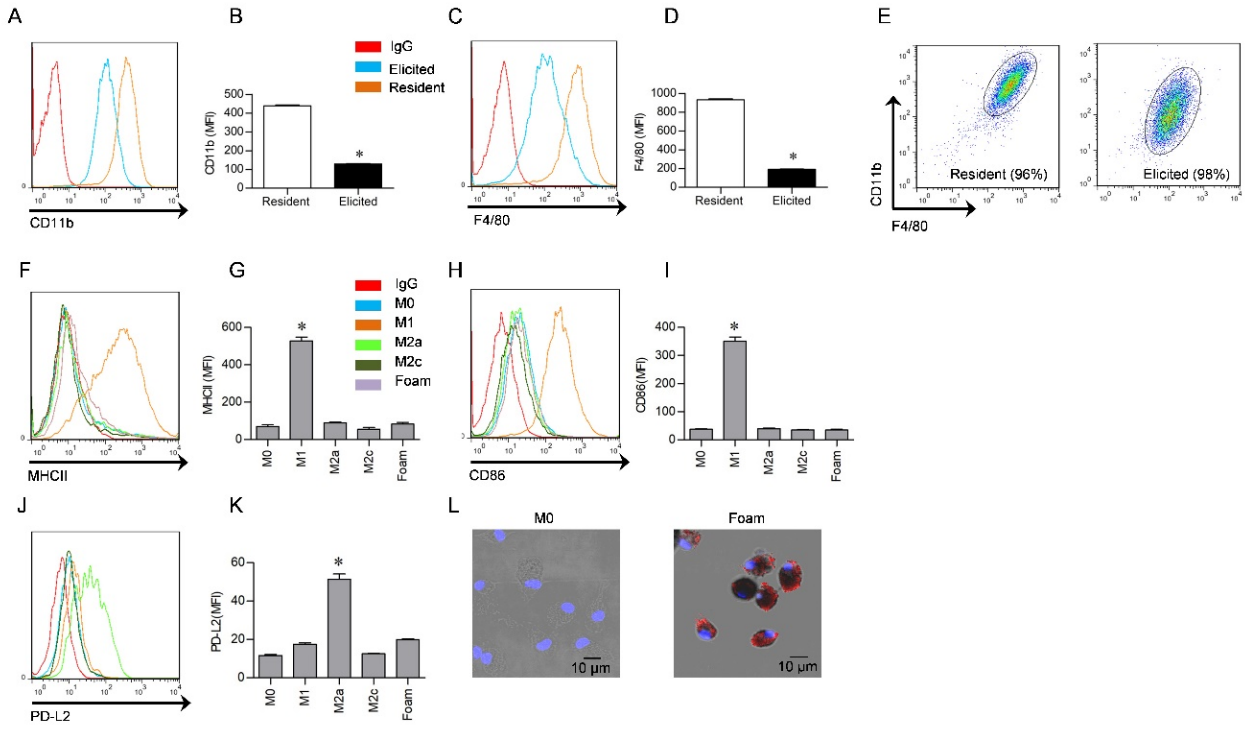


**Figure S1. Characterization of murine peritoneal macrophage subsets.** Resident and elicited macrophages were characterized by expression level of CD11b and F4/80 by flow cytometry. (**A**) Histogram of CD11b. IgG, isotype control. (**B**) Quantification of CD11b mean fluorescence intensity (MFI). (**C**) Histogram of F4/80. (**D**) Quantification of F4/80 MFI. (**E**) The purity of isolated resident and elicited macrophages was assessed by their CD11b and F4/80 levels on scatter plots. (**F-K**) Histogram and MFI from flow cytometry determination of M1 surface markers MHCII and CD86 in addition to M2a surface marker PD-L2. Data are represented as mean ± SEM (n=3). *$P$<0.05 (Student's t-test). (**L**) Micrographs of oil droplet accumulation in foam cells as stained by Oil Red O.

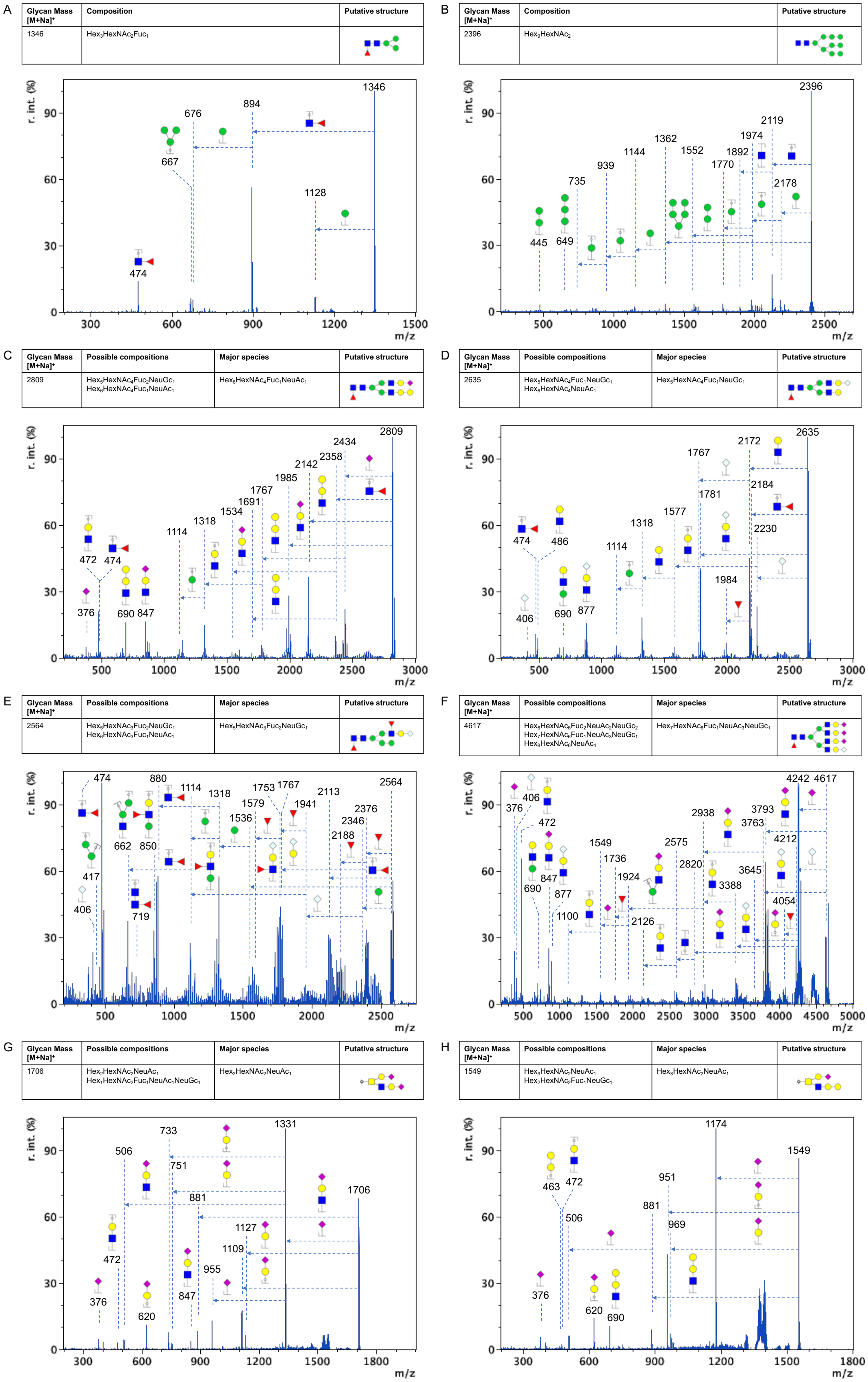


**Figure S2. MALDI-TOF/TOF-MS/MS analysis of permethylated N- and O-glycans obtained from peritoneal macrophage subsets.** Annotations indicate fragment ions from representative (**A-F**) N-glycans and (**G-H**) O-glycans released by β-elimination. Fragmentation patterns were used to derive a putative structure of the major species. Mass spectra were extracted and scaled relative to the maximum intensity (r. int.). Glycan representations follow symbol nomenclature.

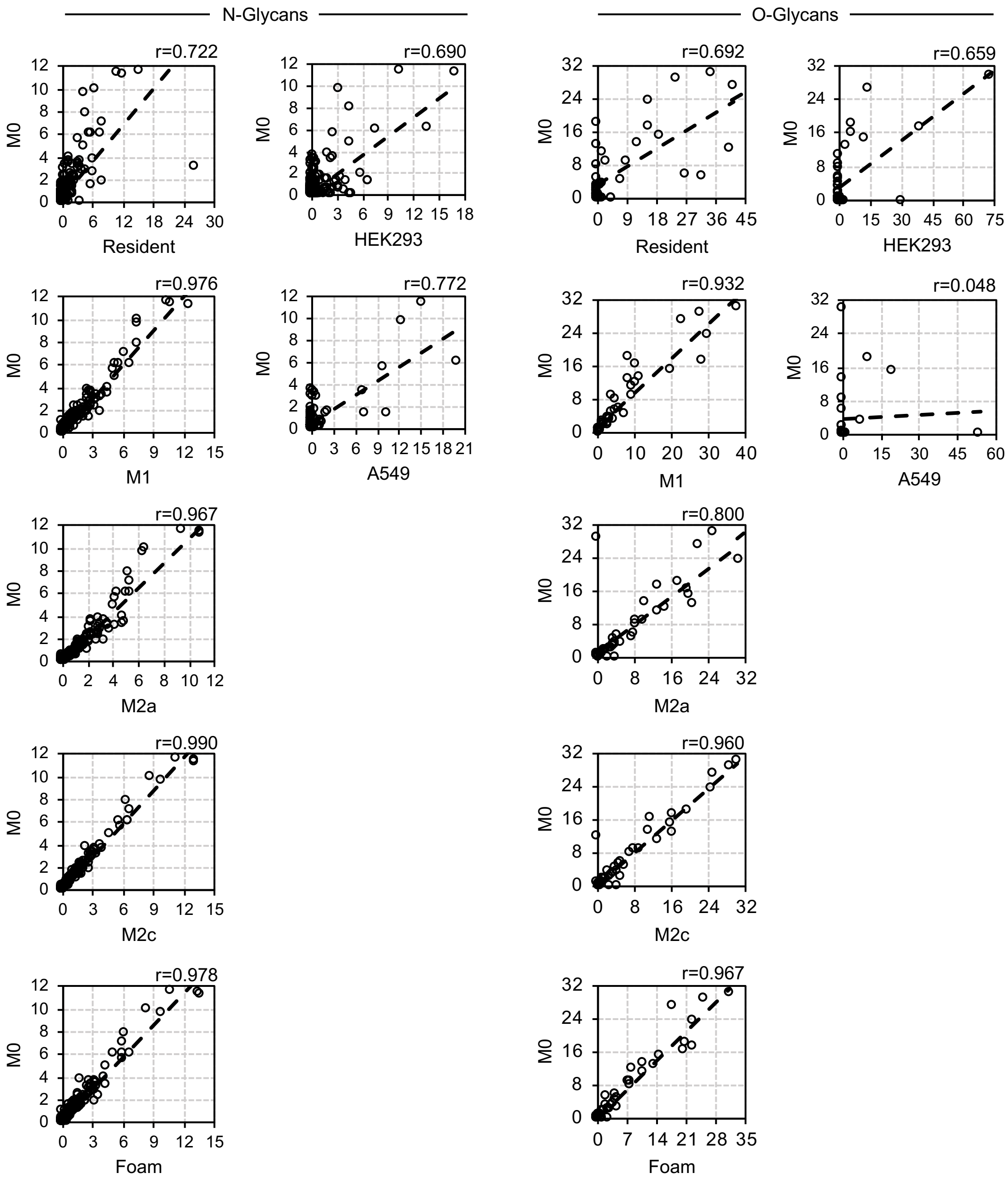


**Figure S3. Correlation scatter plots of N- and O-glycans from M0 vs. resident macrophages, polarized macrophages, HEK293, and A549.** Pearson correlation coefficient values (r) were derived from the average of correlations (n=3/macrophage subset).

A

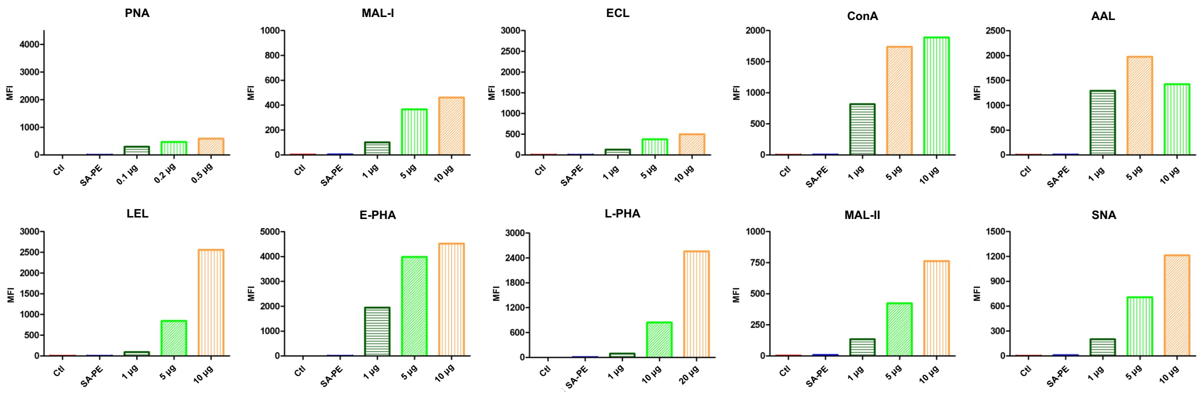


B

| Lectin name | Specificity | Inhibitor |
|---|---|---|
| PNA | asialylated core 1, core 2 | D-galactose |
| MAL-I | Gal-β1,4-GlcNAc | lactose |
| ECL | Gal-β1,4-GlcNAc | lactose |
| ConA | mannose | D-mannopyranose |
| AAL | α1,6-fucose | L-fucose |
| UEA-I | α1,2-fucose | L-fucose |
| LEL | polylactosamine | chitin hydrosylate |
| E-PHA | bisecting GlcNAc | bovine thyroglobulin |
| L-PHA | β1,6-GlcNAc | bovine thyroglobulin |
| MAL-II | α2,3-sialic acid<br>(some additional specificity) | |
| SNA | α2,6-sialic acid | |

C

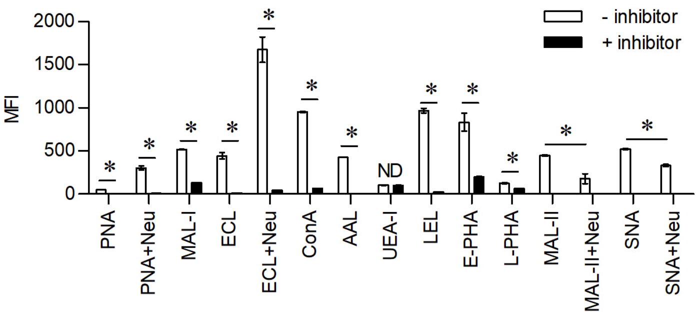


D

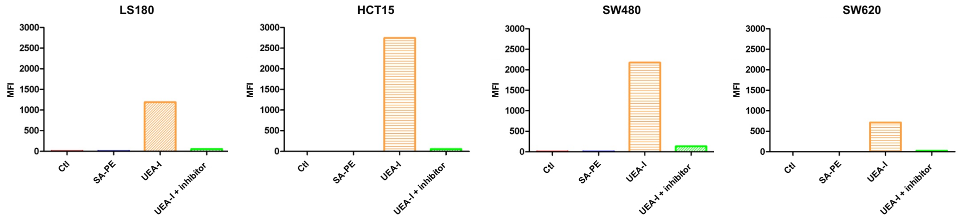


**Figure S4. Optimization of lectin staining against elicited macrophages.** (**A**) Quantification of MFI of bound lectins on elicited macrophages following titration. Ctl, cells only; SA-PE, streptavidin-PE. (**B**) Summary of lectin specificities and inhibitors used in the study. (**C**) Quantification of MFI of bound lectins on elicited macrophages incubated in the presence or absence of inhibitors. For sialic acid binding lectins MAL-II and SNA, a general neuraminidase (Neu) was applied to confirm specificity. Data are represented as mean ± SD (n=2). **P*<0.05 (Student's t-test); ND, not detected. (**D**) UEA-I staining of colon adenocarcinoma cell lines. Positive binding was confirmed by MFI and sequestered by L-fucose, a UEA-I inhibitor.

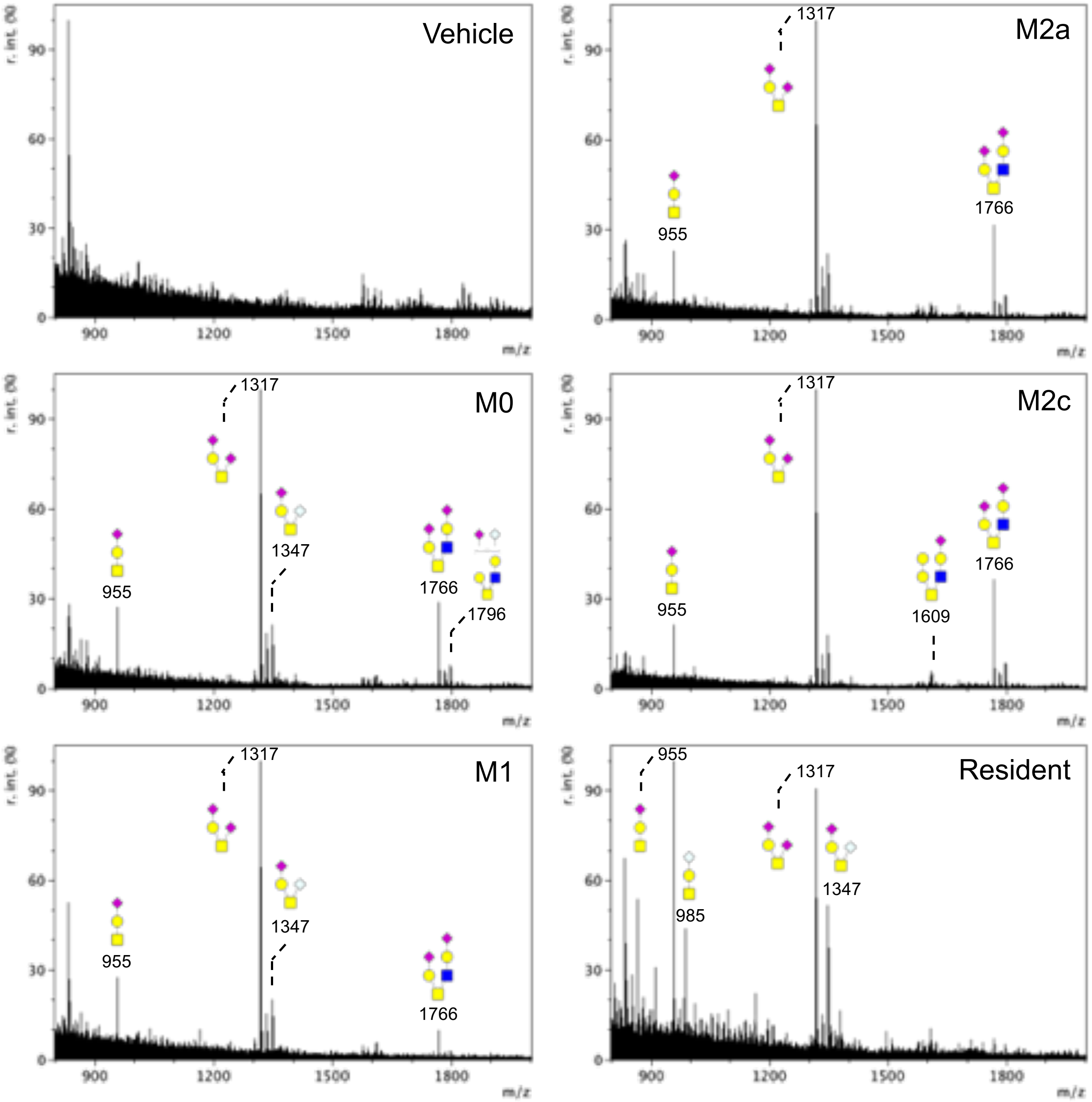


**Figure S5. MS profiles of permethylated O-glycans in peritoneal macrophage subsets obtained from CORA analysis.** Addition of vehicle without $Ac_3$GalNAc-α-Bn was used as a reference control. Mass spectra were extracted and scaled relative to the maximum intensity (r. int.). Glycan representations follow symbol nomenclature.

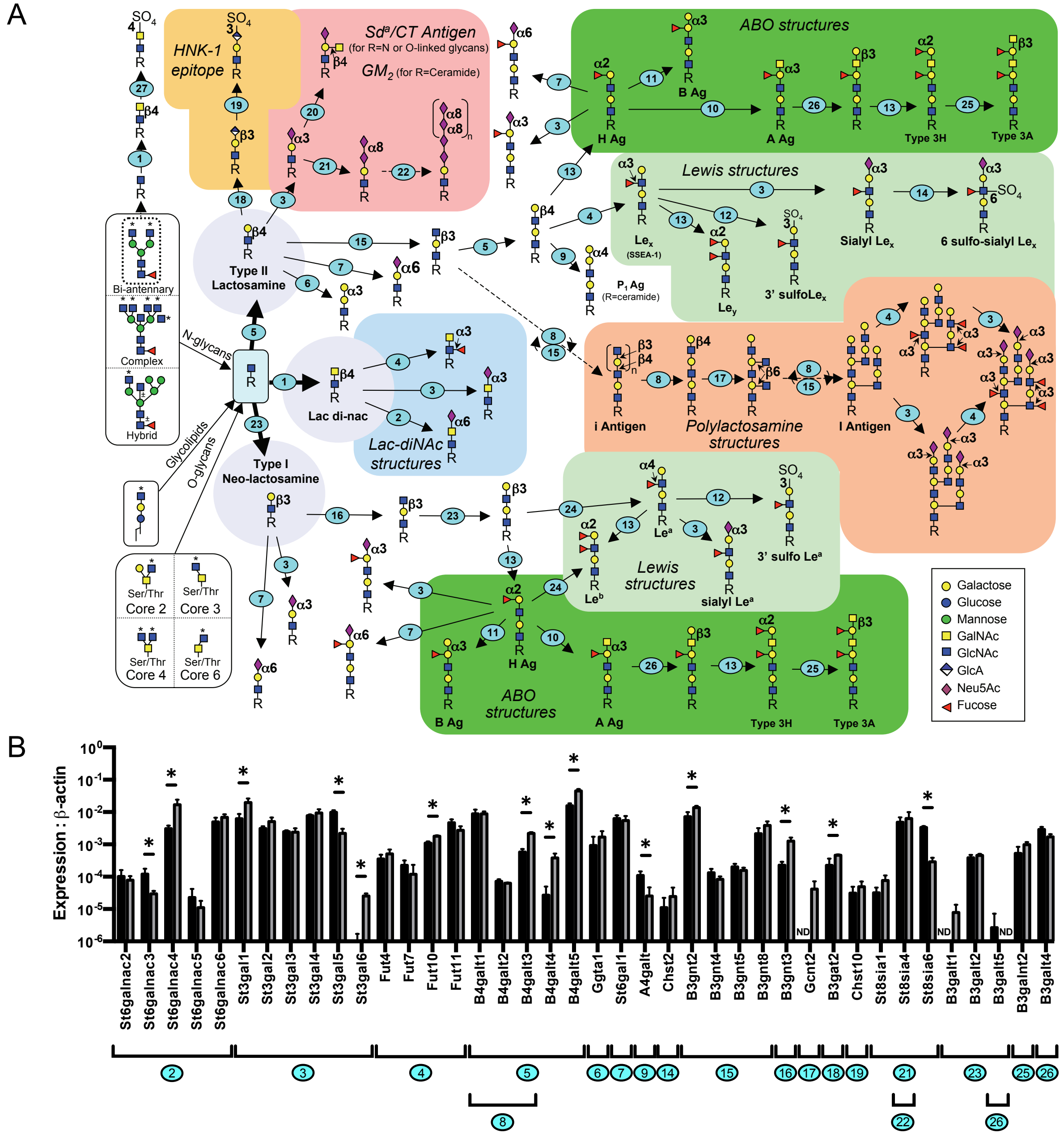


**Figure S6. Quantitative RT-PCR analysis of common genes in the N-glycan, O-glycan, and glycolipid biosynthetic pathways.** Stepwise pathway (**A**) and the associated changes in the expression of genes involved in each step (**B**), which are numbered in sequence. Data were normalized to the expression of β-actin and are represented as mean ± SD (n=4). *$P$<0.05 (Mann-Whitney U test); ND, not detected.

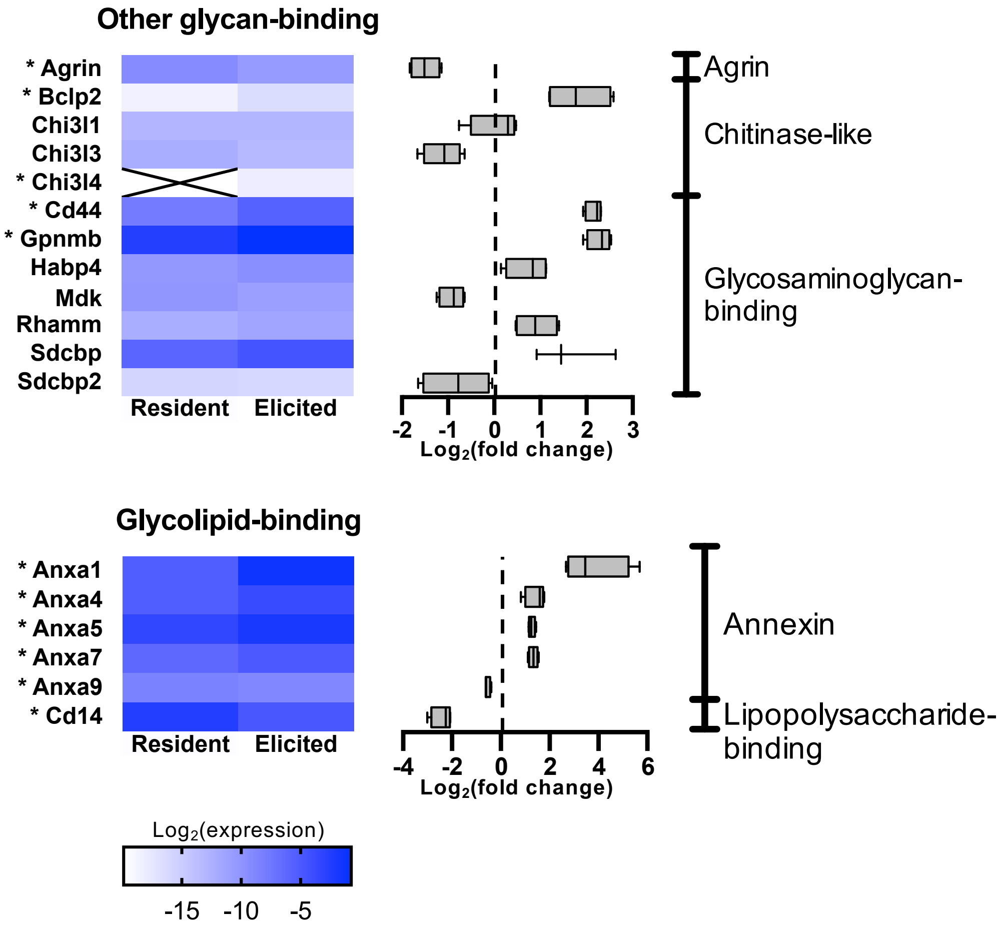


**Figure S7. Quantitative RT-PCR analysis of glycan binding protein genes.** Gene expression in resident and elicited macrophages were quantified and normalized to the expression of β-actin. $Log_2$ transformed mean relative transcript abundances of other glycan and glycolipid-specific protein coding genes (n=4). Box and whisker plots represent fold change of gene expression (resident to elicited). Genes within each indicated family are ordered alphabetically. **P*<0.05 (Mann-Whitney U test); ╳, not detected.
